# Functional networks of the human bromodomain-containing proteins

**DOI:** 10.1101/2022.02.21.481364

**Authors:** Cong Gao, Karen C. Glass, Seth Frietze

## Abstract

**Background:** Bromodomains are a structurally conserved epigenetic reader domain that bind to acetylated lysine residues in both histone and non-histone proteins. Bromodomain-containing proteins (BRD proteins) function are established scaffolds in the assembly of multi-protein complexes to regulate diverse biological processes. BRD proteins have been classified based on biological and functional similarity, however the functions of many BRD proteins remains unknown. PPI network analysis is useful for revealing organizational roles, identifying functional clusters, and predicting function for BRD proteins.

**Results:** We used available data to construct protein-protein interaction networks (PPINs) to study the properties of the human bromodomain protein family. The network properties of the BRD PPIN establishes that the BRD proteins serve as hub proteins that are enriched near the global center to form an inter-connected PPIN. We identified dense subgraphs formed by BRD proteins and find that different BRD proteins share topological similarity and functional associations. We explored the functional relationships through clustering and Hallmark pathway gene set enrichment analysis and identify potential biological roles for different BRD proteins.

**Conclusions:** In our network analysis we confirmed that BRD proteins are conserved central nodes in the human PPI network and function as scaffolds to form distinctive functional clusters. Overall, this study provides detailed insight into the predictive functions of BRD proteins in the context of functional complexes and biological pathways.

## 1 Introduction

Post-translational modifications (PTMs) are fundamental to the dynamic control of protein structure and function. In particular, the acetylation of lysine is an abundant PTM found on both histone and non-histone proteins that is well-known to regulate a variety of biological processes, including transcription, chromatin compaction, protein–protein interactions, cell cycle control, cell metabolism, nuclear transport and actin nucleation (Sterner and Berger 2000, Choudhary, Kumar et al. 2009). Lysine acetylation is reversibly generated by the coordinated actions of both lysine acetyltransferases (KATs) and lysine deacetylases (KDACs) (Shahbazian and Grunstein 2007, Downey and Baetz 2016). Bromodomains are epigenetic reader domains found in a diverse set of chromatin-associated proteins that bind to acetylated lysine residues on histone proteins and non-histone proteins (Fujisawa and Filippakopoulos 2017). The recognition of acetyl-lysine by bromodomain-containing proteins (BRD proteins) and the formation of specific protein-protein and protein-nucleic acid complexes at loci-specific regulatory complexes at functional elements marked by acetyl lysine represents a central mechanism for epigenetic control.

The human proteome encodes 61 BRD domains that are encoded by 42 distinct genes. Each bromodomain is an approximately 110 amino acid structural motif that adopts a 4-alpha-helix barrel structure that forms a binding pocket for acetylated lysine on histones and other proteins (Zeng and Zhou 2002). Most BRD proteins also possess several other conserved functional domains, including other protein-protein interaction or enzymatic domains. Bromodomains can therefore be functionally grouped into 9 distinct classes, including the bromodomain and extra-terminal motif (BET) family, histone modifying factors that either possess intrinsic histone acetyltransferases (HAT), histone methyltransferase (HMT) activities, or belong to subunits of HAT complexes, chromatin remodelling factors, the TRIM/RBCC family, Speckled Proteins (SP), AAA-type ATPase and ZYMND transcriptional repressors (Zaware and Zhou 2019, Boyson, Gao et al. 2021). Although BET proteins and other bromodomain-containing proteins have characterized roles in gene transcription, DNA Damage Repair (DDR) and other chromatin-templated processes, the functions of many bromodomain proteins remain overall poorly described.

The analysis of protein–protein interactions (PPIs) has emerged as a valuable approach to systematically study protein function. High-throughput PPI mapping methodologies including yeast two-hybrid (Y2H) and affinity purification-mass spectrometry (AP-MS) have provided large-scale PPI datasets which are deposited in repositories, including HIPPE (Alanis-Lobato, Andrade-Navarro et al. 2017), HPRD (Peri, Navarro et al. 2003), STRING (von Mering, Huynen et al. 2003), and BIOGRID (Oughtred, Rust et al. 2021). This collection of PPI data makes it possible to create PPI networks (PPINs) to study the network properties based on available graph theory analysis methods that examine static features such as connectivity and location. PPINs can be modelled by undirected graphs, where the nodes are proteins and two nodes are connected by an undirected edge when corresponding proteins physically interact. The representation of PPINs as graphs enables the systematic examination of the topology and function of networks with graph-theoretical principles that can be used to predict the structural properties of the underlying network (Zahiri, Emamjomeh et al. 2020). These predictions provide hypotheses about new interactions from the global network or evidence for exploring functional roles of individual proteins.

To study the role of BRD proteins in the global human interactome, in this study we constructed PPINs based on physical interaction data collected from various resources. We investigated the topological features of the global human PPIN and the sub-network formed by BRD proteins to evaluate the network characteristics of BRD proteins. We further used Hallmark pathway enrichment analysis and clustering with gene ontology to predict the functional characteristics of subnetworks formed by individual BRD proteins. Our results provide confirmation that PPI networks can predict the biological roles of BRD proteins and provide insights on characterizing BRD proteins.

## 2 Materials and Method

### 2.1 Data Description

We collected physical interaction data from the BioGRID (Oughtred, Rust et al. 2021) and HIPPIE (Alanis-Lobato, Andrade-Navarro et al. 2017) databases, then constructed a comprehensive human PPIN (global PPIN), and a sub-network focused on interactions of BRD proteins (BRD PPIN). To evaluate BRD proteins in terms of their relationships, we also constructed a sub-network focused on interactions between BRD proteins (BRD-BRD PPIN).

The BioGRID database collected physical interactions using different techniques from various publications and treats them equally (Oughtred, Rust et al. 2021). We downloaded the version 4.3.194 and extracted all of the physical interactions for homo sapiens. We have summarized the number of different publications or approaches used (shown as the edge width) corresponding to single human protein-protein interactions (**Supplementary Figure 1a**), as well as the interactions for BRD proteins (**Supplementary Figure 1b**). HIPPIE is a specialized human protein-protein interaction network database (Alanis-Lobato, Andrade-Navarro et al. 2017) that integrates interaction data from 10 source databases and 11 studies (that have not been fully covered by the other databases yet), and curates reliable and meaningful interactions via confidence scoring of interactions based on the amount and reliability of evidence supporting them (Alanis-Lobato, Andrade-Navarro et al. 2017). We downloaded the dataset updated in February 2021. The confidence value histogram of physical interactions for BRD proteins and those included in the global PPIN are shown in **Supplementary Figures 2a** and **2b**. Among 391,410 interactions in the global PPIN, 66,711 (1/6) interactions have a confidence value less than 0.63 (suggested confidence value threshold in HIPPIE), and 566 (3,591 in total) interactions for BRD proteins that do not pass the suggested threshold, especially interactions for not well-studied BRD proteins. Therefore, we included all of the human PPI data from BioGRID and HIPPIE and treat them equally without filtering on evidence level. In addition, we also included data from a recent publication (Lambert, Picaud et al. 2019) focused on the interactome of BET proteins. This study revealed non-redundant PPI of BRD2, BRD3 and BRD4 in the native state before addition of JQ1, not yet covered by the HIPPIE or BIOGRID databases. The researchers also used reciprocal methods to explore the interaction between BRD4 and the poorly characterized BRD9 protein and identified new interactors for BRD9 (Lambert, Picaud et al. 2019). The newly identified interactions for BRD2/3/4 and BRD9 are summarized in **Supplementary Figure 3**.

The global PPIN and BRD PPIN are both unweighted and undirected networks without self-loops. We also examined the relationship between the number of publications and BRD protein interaction (degrees) since the representation of interactions in a corresponding PPIN is influenced by the number of publications and the type of study used to report physical interactions (i.e. high throughput assays). Overall, p300 and CBP have the highest number of reported publications (**Supplementary Figure 4**). There seems a positive correlation between the number of publications and degrees in the global and BRD PPINs. Therefore, BRD-BRD PPIN is undirected and weighted by the number of publications and/or techniques, with self-loops indicate the potential formation of homogeneous polymers. Cytoscape (Shannon, Markiel et al. 2003) is used to visualize these networks, BRD proteins are grouped together using group attribute layout (BRD versus non-BRD proteins) and then visualized by degree sorted circle layout.

### 2.2 Graph analysis on the global PPIN and BRD PPIN

The R graph package igraph (Csardi and Nepusz 2006) was used to analyze the topological features of PPINs. Parameters are set to analyze the unweighted and undirected networks. The graph topological features as well as degree, centrality measurements and K-core decomposition for each protein in the global PPIN are computed in this way using the according functions. For graph compactness, if a graph has E ≃ Vk, 2 > k > 1, then this graph is considered as dense, whereas when a graph has E≃V or E≃Vk, k≤1, it is considered as sparse (Koutrouli, Karatzas et al. 2020).

For clique analysis, we aimed to investigate how BRD proteins form functional complexes based the clique prediction. To reduce computation time, we used the according function in igraph package (Csardi and Nepusz 2006) (limiting size >=3) and identify cliques of BRD proteins in BRD PPIN only formed by BRD proteins. The maximal cliques (Eppstein, Löffler et al. 2010) are detected using the function max_cliques in igraph with minimal clique size set to 3. **Table 4** is generated based on the maximal cliques results to show how many BRD proteins in each maximal clique and what they are. We extracted the distance matrix of BRD proteins by applying the distance function in igraph to the global human PPI network and extract the BRD proteins subset. Then the heatmap is plotted using heatmap.2 function in gplots package (Warnes, Bolker et al. 2009). The similarity function in igraph package calculates similarity scores for vertices based on their connection patterns and two nodes can be found to be functionally similar if they share common neighbors (reviewed in (Koutrouli, Karatzas et al. 2020)).

**Table 1.** General topological features of the interaction networks formed by BRD proteins’ interaction profile (BRD PPIN) and all of the human protein-protein interactions (Global PPIN).

|  | BRD PPIN | BRD-BRD PPIN | Global PIN |
| --- | --- | --- | --- |
| Nodes | 4054 | 37/42 | 19,843 |
| Edges | 192,785 | 121 | 559,183 |
| Density | 0.02 | 0.182 | 0.003 |
| Radius | 3 | 2 | 4 |
| Diameter | 4 | 4 | 8 |
| Average shortest paths | 2.21 | 2.123 | 2.87 |
| Average clustering coefficients | 0.22 | 0.475 | 0.14 |
| Connected component | 1 | 6 | 18 |

**Table 2.** Statistical summary of centrality and K-core measurements for BRD proteins vs. non-BRD proteins in the global PPIN. The significance level is 0.05.

|  | BRD proteins | Non-BRD proteins | P value (two-sided T test) |
| --- | --- | --- | --- |
| Degree mean | 207.93 | 56.08 | $2.210 \times 10^{-16}$, * |
| Betweenness Centrality mean | 97567.05 | 18340.94 | 0.026, * |
| Closeness Centrality mean | $2.24 \times 10^{-6}$ | $2.20 \times 10^{-6}$ | $5.78 \times 10^{-14}$, * |
| Eccentricity mean | 5.048 | 5.387 | $7.638 \times 10^{-13}$, * |
| Eigenvector Centrality mean | 0.103 | 0.028 | 0.00103, * |
| Clustering coefficients mean | 0.133 | 0.141 | 0.6172, *ns* |
| K-core | 60.02 | 28.75 | $1.685 \times 10^{-13}$, * |

**Table 3.** Maximal cliques in the interaction profile between BRD proteins

| **Clique Index** | **BRDs** | **Number of nodes** |
| --- | --- | --- |
| 1 | BRDT, BRD2, BRD4 | 3 |
| 2 | BRD1, PBRM1, TRIM33 | 3 |
| 3 | BPTF, BRD2, BRD3, BRD4 | 4 |
| 4 | ZMYND11, BRD4, SMARCA2 | 3 |
| 5 | ZMYND11, BRD4, BRPF3 | 3 |
| 6 | BAZ2A, BAZ1B, BRD2 | 3 |
| 7 | BAZ2A, BAZ1B, TRIM33 | 3 |
| 8 | KMT2A, SMARCA2, BRD4, TAF1 | 4 |
| 9 | KMT2A, SMARCA2, BRD4, CBP | 4 |
| 10 | BRPF1, BRPF3, TAF1 | 3 |
| 11 | BRPF1, BRD2, TAF1 | 3 |
| 12 | BRPF1, BRD2, ZMYND8 | 3 |
| 13 | BRPF1, BRD2, TRIM24 | 3 |
| 14 | KAT2B, CBP, P300, SMARCA2 | 4 |
| 15 | KAT2B, CBP, P300, KAT2A | 4 |
| 16 | KAT2A, CBP, BRD4, P300 | 4 |
| 17 | P300, SMARCA2, BRD7 | 3 |
| 18 | P300, SMARCA2, BRD4, CBP | 4 |
| 19 | BAZ1A, BRD4, BRD3, SMARCA2 | 4 |
| 20 | BAZ1A, BRD4, BRD3, BAZ1B | 4 |
| 21 | BRD8, TAF1, BRPF3, SP110, BRD4 | 5 |
| 22 | ZMYND8, SMARCA4, BRD4, TRIM28 | 4 |
| 23 | ZMYND8, SMARCA4, BRD4, BRD3, BRD2 | 5 |
| 24 | CBP, TRIM28, TRIM24 | 3 |
| 25 | CBP, TRIM28, BRD4, SMARCA4, SMARCA2 | 5 |
| 26 | PHIP, BRD2, BRD3, BRD4, TAF1 | 5 |
| 27 | PHIP, BRD2, BRD3, BRD4, PBRM1, SMARCA4 | 6 |
| 28 | TRIM24, BRD2, BRD7 | 3 |
| 29 | TRIM24, TRIM28, BRD7 | 3 |
| 30 | TRIM24, TRIM28, TRIM33 | 3 |
| 31 | TAF1, SMARCA2, BRD3, BRD4, BRD2 | 5 |
| 32 | BRD9, SMARCA2, BRD3, BRD4, BRD2, SMARCA4 | 6 |
| 33 | PBRM1, SMARCA2, SMARCA4, TRIM33, BRD4 | 5 |
| 34 | PBRM1, SMARCA2, SMARCA4, BRD3, BRD2, BRD7 | 6 |
| 35 | PBRM1, SMARCA2, SMARCA4, BRD3, BRD2, BRD4 | 6 |
| 36 | SMARCA2, SMARCA4, TRIM28, BRD7 | 4 |
| 37 | SMARCA2, SMARCA4, TRIM28, TRIM33, BRD4 | 5 |
| 38 | SMARCA4, BAZ1B, BRD4, TRIM28, TRIM33 | 5 |
| 39 | SMARCA4, BAZ1B, BRD4, BRD3, BRD2 | 5 |

**Table 4.**
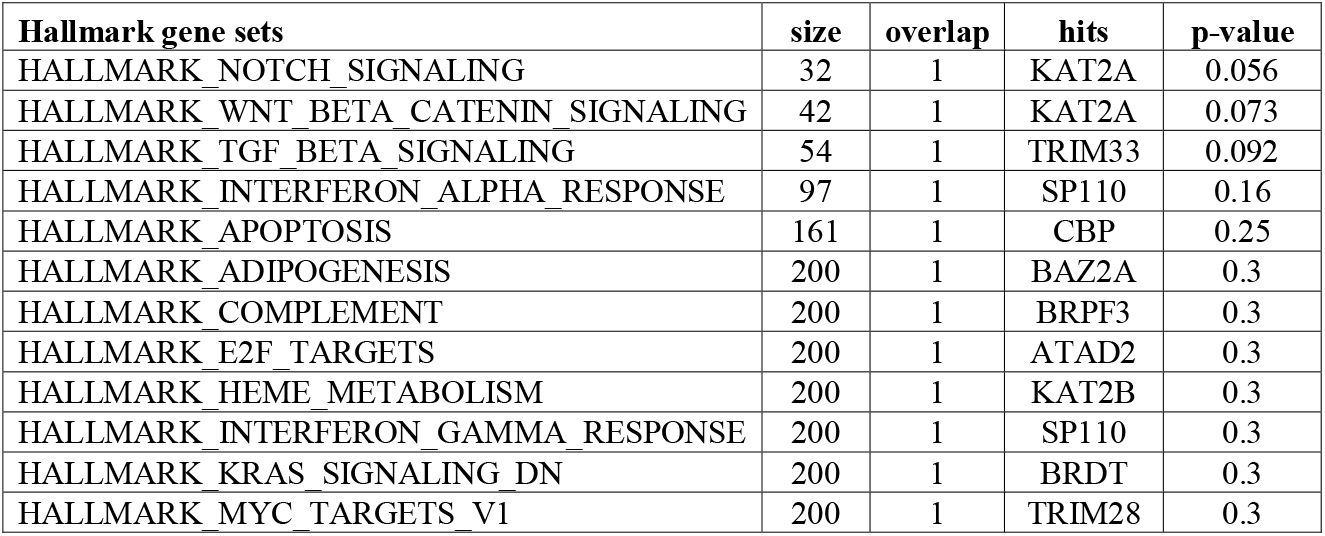
Involvement of 42 BRD proteins in Hallmark pathway gene sets

| Hallmark gene sets | size | overlap | hits | p-value |
| --- | --- | --- | --- | --- |
| HALLMARK_NOTCH_SIGNALING | 32 | 1 | KAT2A | 0.056 |
| HALLMARK_WNT_BETA_CATENIN_SIGNALING | 42 | 1 | KAT2A | 0.073 |
| HALLMARK_TGF_BETA_SIGNALING | 54 | 1 | TRIM33 | 0.092 |
| HALLMARK_INTERFERON_ALPHA_RESPONSE | 97 | 1 | SP110 | 0.16 |
| HALLMARK_APOPTOSIS | 161 | 1 | CBP | 0.25 |
| HALLMARK_ADIPOGENESIS | 200 | 1 | BAZ2A | 0.3 |
| HALLMARK_COMPLEMENT | 200 | 1 | BRPF3 | 0.3 |
| HALLMARK_E2F_TARGETS | 200 | 1 | ATAD2 | 0.3 |
| HALLMARK_HEME_METABOLISM | 200 | 1 | KAT2B | 0.3 |
| HALLMARK_INTERFERON_GAMMA_RESPONSE | 200 | 1 | SP110 | 0.3 |
| HALLMARK_KRAS_SIGNALING_DN | 200 | 1 | BRDT | 0.3 |
| HALLMARK_MYC_TARGETS_V1 | 200 | 1 | TRIM28 | 0.3 |

To extend the similarity measurements beyond the direct neighbors of each node, we analyzed the distance matrix from the global PPIN and obtained the indirect interaction partners list for each BRD proteins in the global human PPI network if the partners are away from the BRD proteins with the shortest path of 2 and 3 (denotes as SP=2 and SP=3), respectively. Then we use the modified mathematical definition:

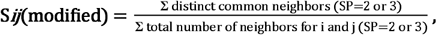

i and j denote different BRD proteins to calculate the extended similarity of each pair of BRD proteins and compare the indirect interaction profiles of two BRD proteins.

### 2.3 Statistical Analysis

Direct comparisons of topological features between BRD proteins and non-BRD proteins were performed by two-tailed un-paired Student T test (Student 1908). Chi-Square test is performed based on degree mean in the global PPIN. We used the fit_power_law function in igraph package to check whether we can fit a power-law distribution to the degree distribution in the global human PPI network (Csardi and Nepusz 2006). The ‘plfit’ implementation is used for this function attempting to find the optimal value of the fitted power law distribution for which the p-value of a Kolmogorov-Smirnov test between the fitted distribution and the original sample is the largest. Under this setting, we checked the p-values that is based on the hypothesis that the original data could have been drawn from the fitted power-law distribution.

### 2.4 Pathway enrichment analysis and functional clustering

Pathway enrichment analysis was performed on BRD proteins and their non-BRD interaction partners using hypeR bioconductor package (Federico and Monti 2020). Hallmark gene sets are obtained from the Molecular Signatures Database v7.4 (MSigDB) (Liberzon, Birger et al. 2015). For the pathway enrichment on non-BRD interacting proteins, we extracted the top 25 Hallmark pathways which are sorted by FDR (<0.05) and the overlapped non-BRD proteins for each top enriched pathway. Then we generated interaction profile for each BRD proteins in each top enriched pathway by computing the number of interacting non-BRD proteins for the specific BRD proteins and also participating in the specific biological pathway. For individual pathway subnetworks, we extracted the interactions of BRD proteins involved in DNA repair, MTORC1 signaling pathway and oxidative phosphorylation process, then used Cytoscape (Shannon, Markiel et al. 2003) to visualize the pathway-focused interaction networks. MTGO was used to combine graph topology and gene ontology (Vella, Marini et al. 2018). We obtained the gene annotation and files from Gene Ontology (GO) database (Ashburner, Ball et al. 2000) and extracted the GO term file from Go.db package (Carlson, Falcon et al. 2017) in R. MTGO generates topological modules denoted as set G based on the graph topology, and functional modules, set T, in which each set member correspondent to one GO term. We extracted the GO IDs and gene symbols corresponding to each protein as input for MTGO using minSize of 8 and maxSize of 300. Then we combine the optimized clustering file with GO description and extracted the clusters in which BRD proteins are the cluster members.

## 3 Results

### 3.1 Construction of a bromodomain protein interaction network

To study the global interaction properties of BRD proteins, we constructed a network of the physical interactions of all 42 members of the bromodomain protein family (see methods). The bromodomain protein family protein-protein interaction network (BRD PPIN) is comprised of 4,054 unique interacting proteins (nodes) and 192,785 non-redundant edges (**Table 1**). In comparison, the complete human PPIN (global PPIN) contains a total of 19,843 nodes and 559,183 edges.

Among BRD proteins, TRIM28, BRD4, CBP and p300 have the highest number of interactions, whereas SP140, SP140L and ATAD2B have the lowest number of interactions in the BRD PPIN (**Figure 1**). Combined with the publication numbers for each BRD protein, BRD1, BRD2, BRD3, BRD4, BRD7 and TRIM28 have a higher number of degrees, even with a relatively smaller number of publications compared to its highest degree. SMARCA4, KAT2B and p300 show a consistent number of degrees compared to the number of publications, whereas SMARCA2, BRD2 and TRIM28 have higher degrees and moderate number of publications.

**Figure 1.**
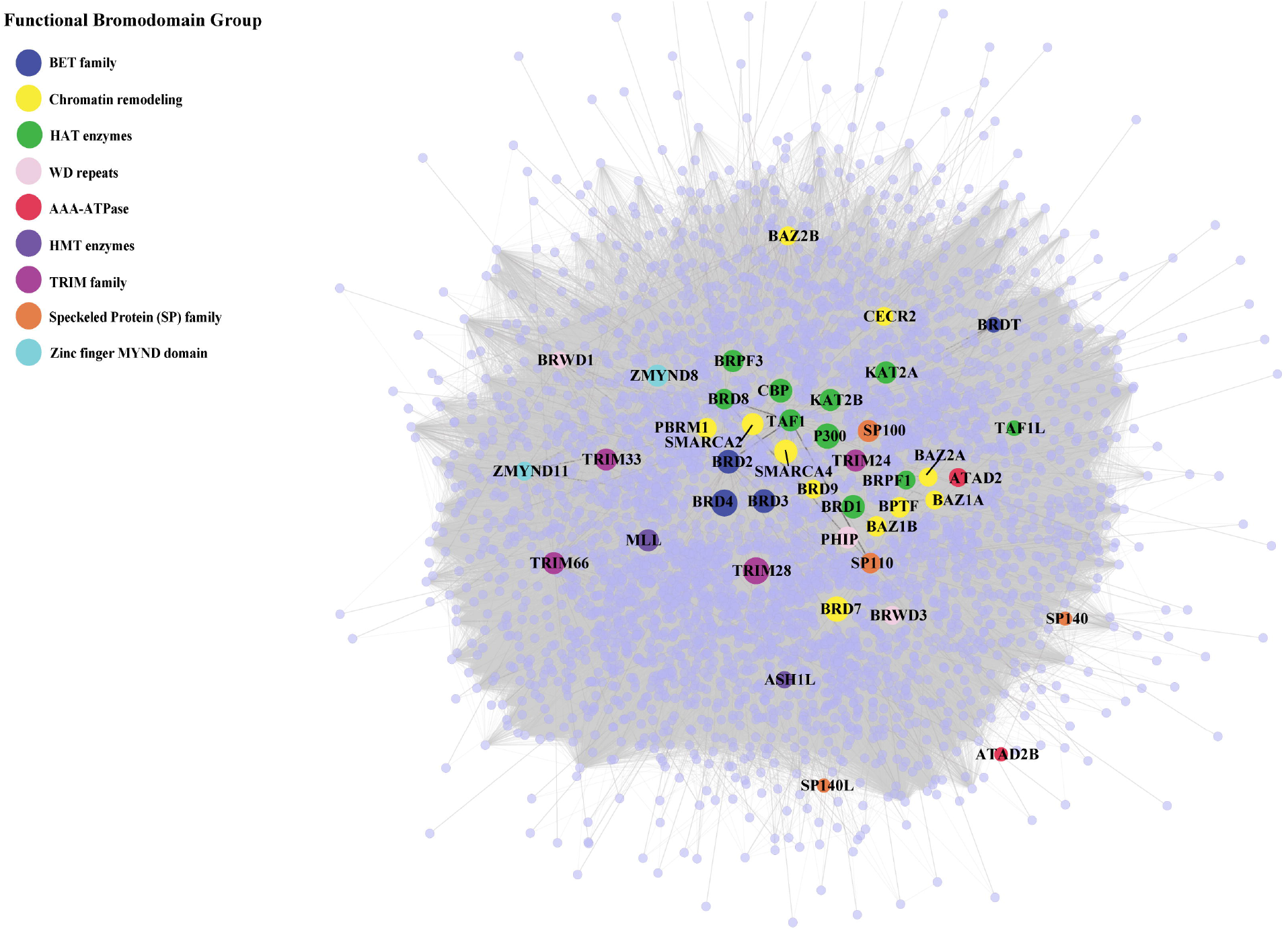

### 3.2 The BRD PPIN is a connected scale-free network

Protein interaction networks exhibit specific topological features that depict the biological properties of the network. The global PPIN can be divided into 18 distinct components based on the connected components analysis and eccentricity, whereas the BRD PPIN forms a single connected component (**Table 1**). The BRD PPIN exhibited a smaller radius, diameter and average shortest path compared to the global PPIN, suggesting that BRD proteins form a tighter network and are inter-connected. The edge density of BRD PPIN is 0.02, indicating that only 2% of the total possible number of edges are observed, possibly due to the fact that many physical interactions remain to be discovered, especially for less studied BRD proteins. The clustering coefficient (cliques, or the formation of complete subnetworks) is a measure of whether a node has the tendency to form clusters or tightly connected communities (e.g., protein clusters in a protein-protein interaction network) (Koutrouli, Karatzas et al. 2020). While the probability of forming cliques consisting of three or more nodes is low in the BRD PPIN, it is considerably higher than that of the global PPIN (clustering coefficient 0.14 versus 0.22, respectively). The degree distribution of nodes and the cumulative frequency curve in BRD PPIN and the global PPIN indicates that a small number of nodes have high degrees (**Figure 2**). The degree distribution suggests these are scale-free networks. A biological network that is scale-free is stable and tolerant to perturbations and is also venerable to loss of hub proteins in the overall networks. We further confirmed that BRD PPIN and the global PPIN are both scale-free networks by carrying out Kolmogorov-Smirnov (KS) statistical test (p-value=0.91 and KS=0.022 for the global PPIN; p-value=0.643 and KS=0.033 for BRD PPIN). Taken together, these results indicate that BRD proteins form a densely connected network with their interaction partners.

**Figure 2.**
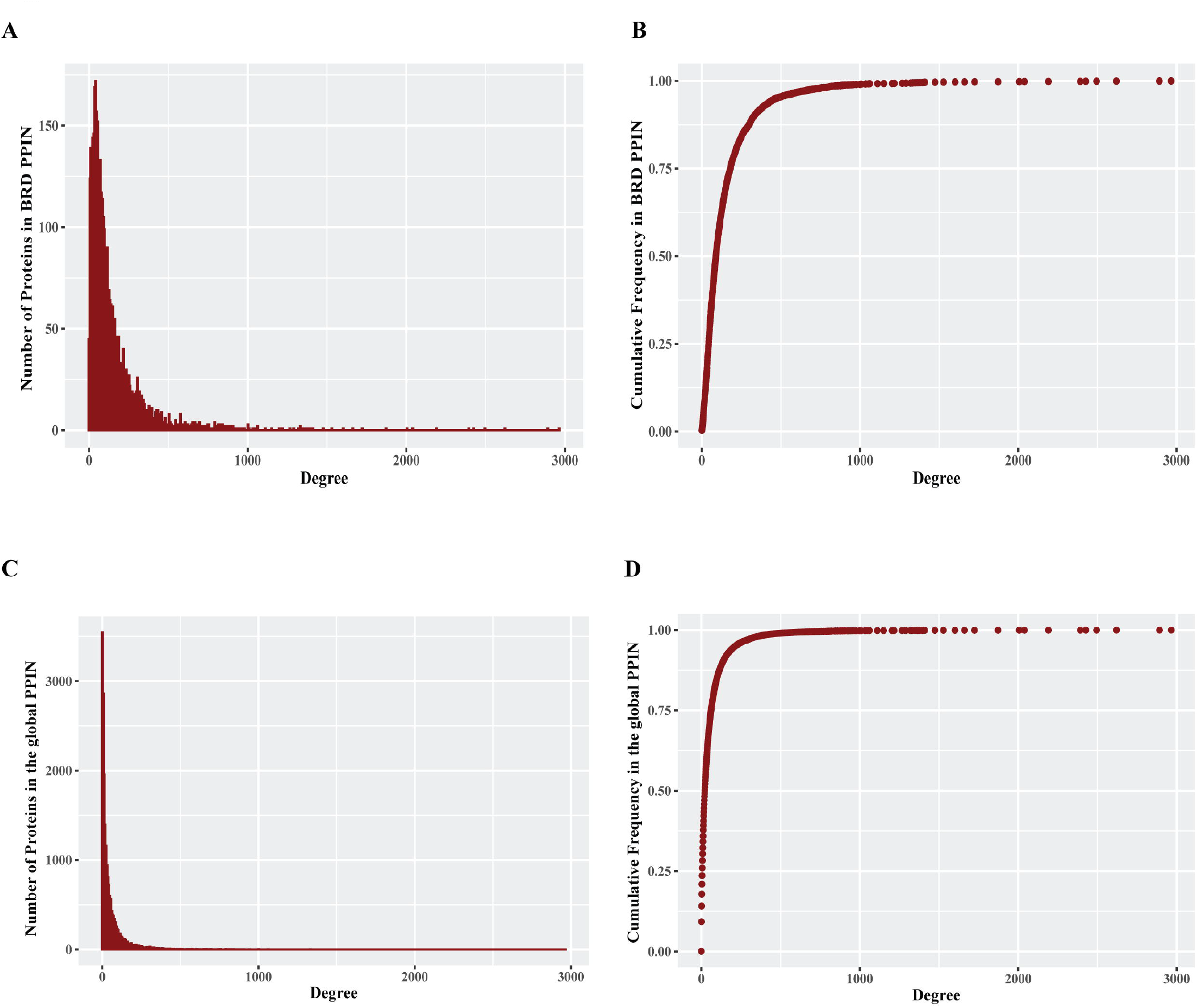

### 3.3 Bromodomain proteins serve as central nodes in the global human PPI network

We next focused on BRD proteins and determined specific topological properties for each node including degree, centrality measurements and K-core decomposition. We investigated the role of BRD proteins as hubs in the PPIN. Hub proteins are proteins that form a high number of interactions in the network and are important for the formation of PPI clusters (Barabasi 2012). We assessed whether BRD proteins have more interactions than non-BRD proteins. The mean degree of BRD proteins is 207.93, compared to 56.08 for non-BRD proteins (**Table 2**, p-value<0.01). We calculated the mean degree of all nodes in the global PPIN and defined proteins with a higher degree than the mean degree as hub proteins. We then compared the percentage of hub proteins between BRD protein family to non-BRD protein family. This analysis showed that BRD proteins are more likely to serve as hubs in the entire human network (Chi-Square test, p-value<0.01; see methods). BRD2, BRD3, BRD4, CBP, SMARCA2 and SMARCA4 are among the top hub BRD proteins (**Figure 1 & Supplementary Figure 4**).

To further evaluate the importance of BRD proteins in the robustness of the network (tolerance to perturbations) and their role in bridging different

PPI communities, we examined other centrality measurements of BRD proteins in the global PPIN. Statistical inference indicates that BRD proteins have a significantly higher betweenness centrality mean than that of non-BRD proteins (**Table 2**, p-value<0.05). The eigenvector centrality is a measure of the influence of a node in a network, and higher eigenvector centrality scores indicate this node is connected to other high influential nodes (Chen, Liao et al. 2019, Koutrouli, Karatzas et al. 2020). BRD proteins exhibit a significantly higher closeness centrality mean and smaller eccentricity mean as well as higher eigenvector centrality mean (**Table 2**, p-value<0.05). These results suggest that BRD proteins are influential members in the global PPIN.

Proteins are hierarchically located in the PPIN, and those with high degrees are more likely to be located near or at the center of the entire network (Wuchty and Almaas 2005). However, the degree numbers alone do not reflect the hierarchies and the locations of proteins within the network (Jeong, Mason et al. 2001, Zou, An et al. 2018). We therefore performed K-core decomposition process to split the network into different layers from outside to inside in order to understand BRD protein functions in the network organization (Alvarez-Hamelin, Dall’Asta et al. 2005). Hub nodes with higher K-core values are referred as a global center in the whole network, and hubs with relatively lower K-core values are the local centers to forming the periphery connected clusters. The global PPIN is split into 102 layers and the K-core for BRD proteins ranges from 4 to 102. As the percentage of BRD proteins in most layers keep zero since there are many layers not including BRD proteins, there is no clear relationship between BRD proteins percentage in each layer and K-core numbers. We compared the K-core means between BRD proteins and non-BRD proteins and found that the K-core mean for BRD proteins are significantly higher than that of non-BRD proteins (3-fold over non-BRD proteins, **Table 2**). In the global PPI network, BRD proteins are more likely to be located near the global center of the network topological organization. As it has been demonstrated that proteins near the global center in the yeast PPIN tend to be essential and conserved in evolution (Wuchty and Almaas 2005), this information also provides support that BRD proteins tend to be essential and conserved in evolution across different organisms. Among BRD proteins, BRD4, TRIM28, p300 and BRD7 have the highest K-core values with highest degrees. Taken together the result of this topological analysis indicates BRD proteins perform important organizational functions for the global PPIN and their roles may be evolutionary conserved.

### 3.4 Bromodomain-containing proteins cooperate and exhibit functional similarities

Connected proteins within the PPIN may share similar functions and studying the relationships between BRD proteins will be helpful to predict the function of less-well characterized BRD proteins. To investigate the interactions among BRD protein family members we constructed a subnetwork based on the interactions between BRD proteins (BRD-BRD PPIN). Several BRD proteins exhibited no less than 10 non-redundant interactions with other family members, including BRD2, BRD3, BRD4, SMARCA2, SMARCA4, TAF1, CBP, and TRIM33 (**Figure 3**). Self-loops measure the potential of proteins to form dimers or oligomers. For example, ATAD2 subunits has been reported to form hexamers (Cho et al., 2019). Among BRD proteins, 17 members form self-loops (**Figure 3**). Among these, the self-loop of p300 has the largest width, and this protein has also been reported to homo-oligomerize (Zhang et al., 2021). The self-loops do not seem related to degrees in the entire human network, or in BRD-BRD PPIN.

**Figure 3.**
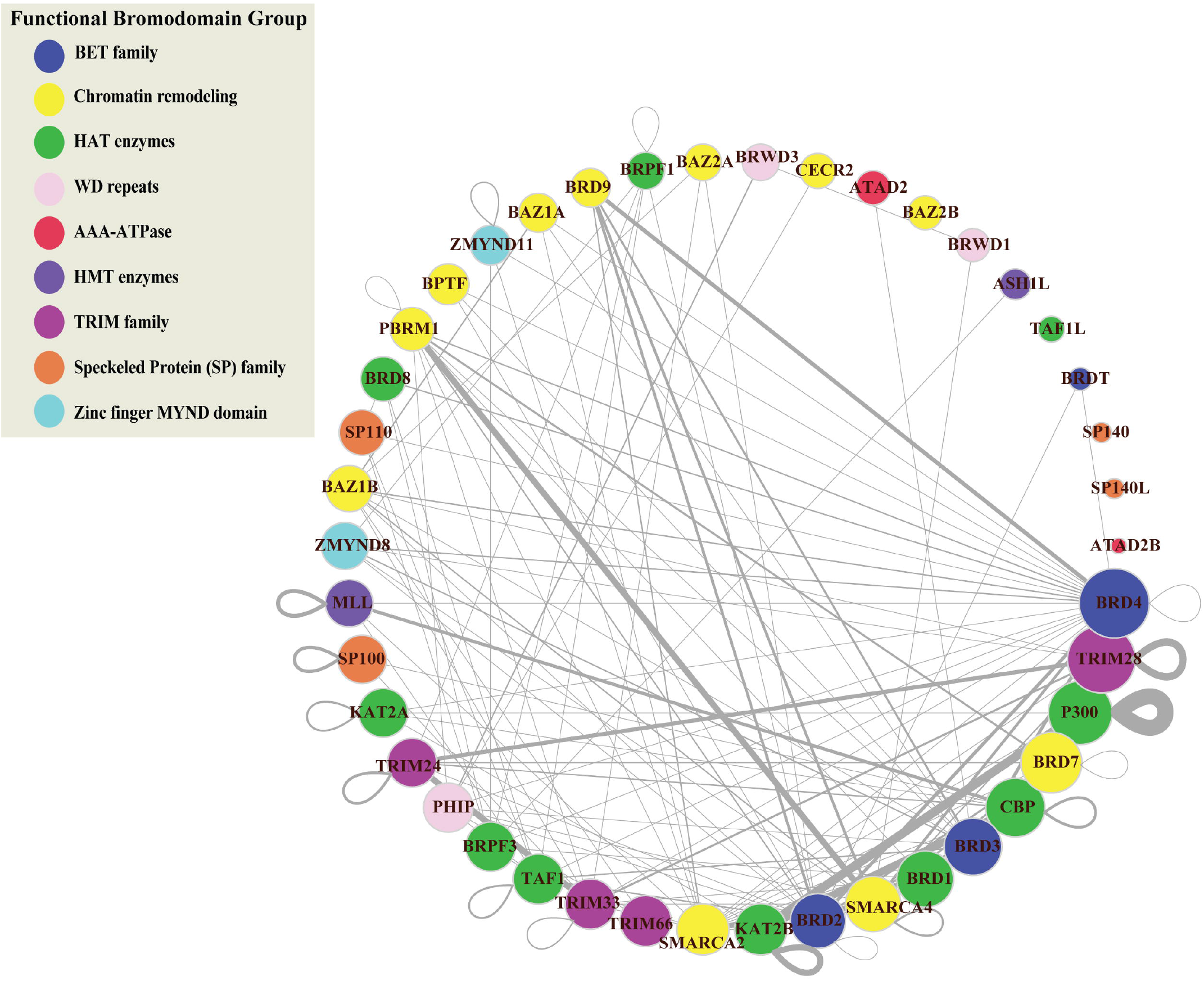

Cliques in PPI networks are related to protein complexes and functional modules that have a biological significance (Zahiri, Bozorgmehr, & Masoudi-Nejad, 2013), and components in protein complexes or functional modules are prone to interacting with each other (Zahiri et al., 2013). We performed clique detection on the BRD-BRD PPIN (Zahiri et al., 2013). We identified 273 cliques formed by BRD proteins with size greater or equal to 3 nodes. Smaller cliques can be merged to form maximal cliques that cannot be extended by including one more adjacent vertex. In total, we identified 39 maximal cliques in the BRD-BRD PPIN (**Table 3**). We also used shortest path and similarity measurements from the global human PPI network to the relationship between BRD proteins. Hierarchical clustering of the shortest paths for each BRD protein reveals a cluster of BRD proteins that tend to interact with each other or form complexes (**Figure 4A**). The shortest paths between different BRD proteins ranged from 1 to 3. Most BRD protein pairs exhibited a shortest path of 2 and are connected by another protein (BRD protein or non-BRD protein). For example, there is no direct link between TRIM66 and ATAD2B, but they are connected by one non-BRD interacting proteins (**Figure 4B**). These shorter distances, like the “six degrees of separation” concept, indicates that the cellular interactome between BRD proteins is relatively small, and suggests BRD proteins tend to be connected closely together allowing for fast communication with each other.

**Figure 4.**
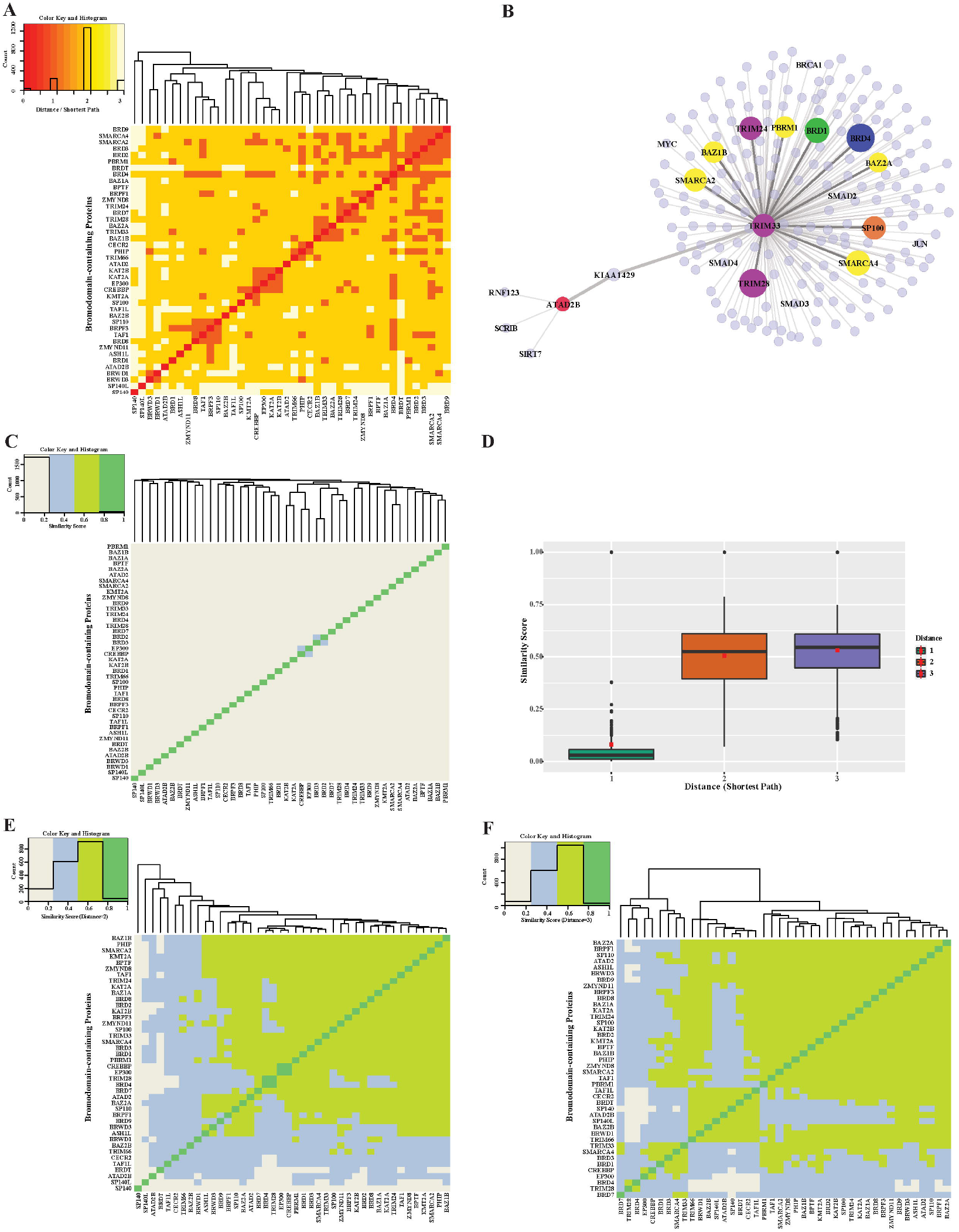

The close connection of BRD proteins is a result of the PPIN topology and likely their functional relationships. The similarity interaction heatmap shows the similarity scores of adjacent neighbors for each pair of BRD proteins (**Figure 4C**). Most BRD protein pairs harbor low similarity values. There are 4 pairs of nodes with relatively higher similarity values including, CBP and P300, SMARCA2 and SMARCA4, KAT2A and KAT2B, and BRD2 and BRD3. This similarity score measurement only accounts for neighboring interactions (shortest path = 1) but failed to reveal similarities beyond these pairs of proteins, which share structural similarity, and belong to similar functional groups. We therefore modified the similarity score to show the ratio of common indirect interacting partners with distances of 2 and 3 (denoted as modified similarity). The similarity score within different BRD proteins increases dramatically after extending the shortest paths to 2 or 3 and the mean similarity score increases by about five-fold (**Figure 4D**). Most pairs of BRD proteins have a modified similarity score greater than 0.5 (**Figures 4E and 4F**).

### 3.5 Bromodomain-containing proteins are enriched in various Hallmark and Gene Ontology pathways

Several BRD proteins have been functionally characterized or classified into functional groups based on the presences of conserved protein domains outside of the bromodomain.

Approximately one-quarter of all BRD proteins (10/42) are curated as members of Hallmark pathways from Molecular Signatures Database (MSigDB) (Liberzon et al., 2015) (**Table 4**). Among these, only KAT2A and SP110 belong to more than one Hallmark pathway. How-ever, without available data for the less well-studied BRD proteins a systematic analysis of PPIs is needed to predict their functional roles. We therefore performed pathway enrichment analysis using the 4,012 non-BRD interacting proteins in the BRD PPIN to examine the global roles of BRD proteins. Among the 50 Hallmark pathways, 42 of have a significance level less than 0.05 (FDR, false discovery rate), indicating these non-BRD interactors are significantly enriched in a variety of Hallmark pathways (**Figure 5A**). The gene sets of transcriptional fac-tors, such MYC and E2F targets, as well DNA repair and cell cycle checkpoints gene sets are among the most highly enriched pathways. Interestingly, many BRD members are associated with a given pathway. For example, a total of 32 BRD proteins are involved in DNA repair response via interaction with members of this pathway (**Figure 5B**).

**Figure 5.**
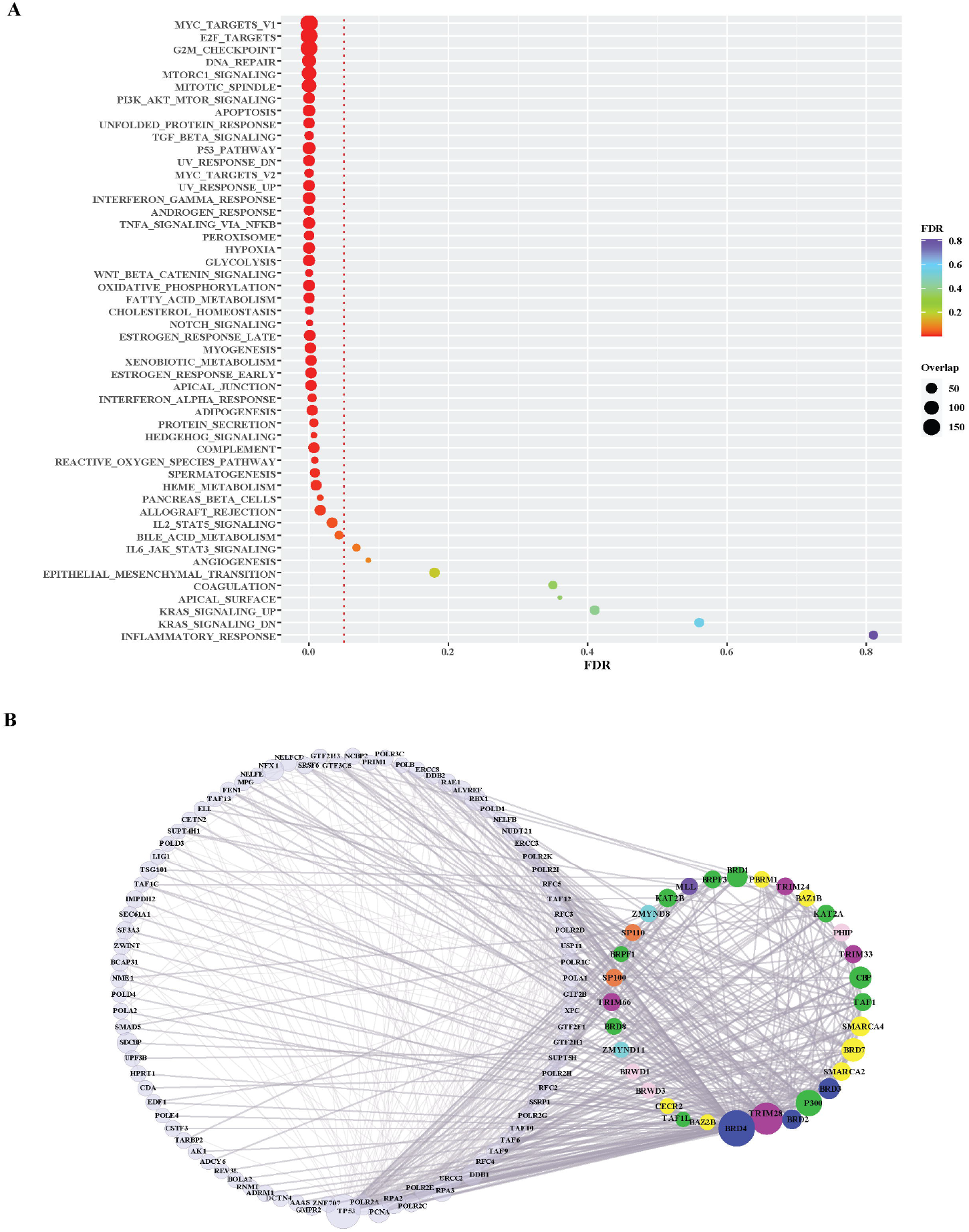

We further investigated the level of each BRD protein involved into the top 25 significantly enriched Hallmark pathways. Overall, the inter-action partners of BRD4 and TRIM28 are the most enriched BRD proteins among these different Hallmark pathways (**Figure 6A**). BRD4 interactions are highly enriched in mTORC1 signaling and glycolysis (**Figure 6B**). The inhibitory function of the mTOR complex 1 (mTORC1) in autophagy is well established (reviewed in (Jung, Ro, Cao, Otto, & Kim, 2010)) and BRD4 has been characterized as a transcriptional repressor of autophagy and lysosomal function (Sakamaki et al., 2017). The function of BRD4 as gene transcriptional regulator to modulate glycolysis has been studied (Xu et al., 2021). All these findings give support to the importance of BRD4 in mTORC1 signaling and glycolysis. A sub-network focus on the interaction between mTORC1 pathway members and BRD proteins show that 28 BRD proteins also interact with this pathway. These results indicate that a large proportion of BRD proteins potentially play roles in the mTORC1 signalling pathway, but the exact mechanistic roles of these BRD proteins are yet to be discovered.

**Figure 6.**
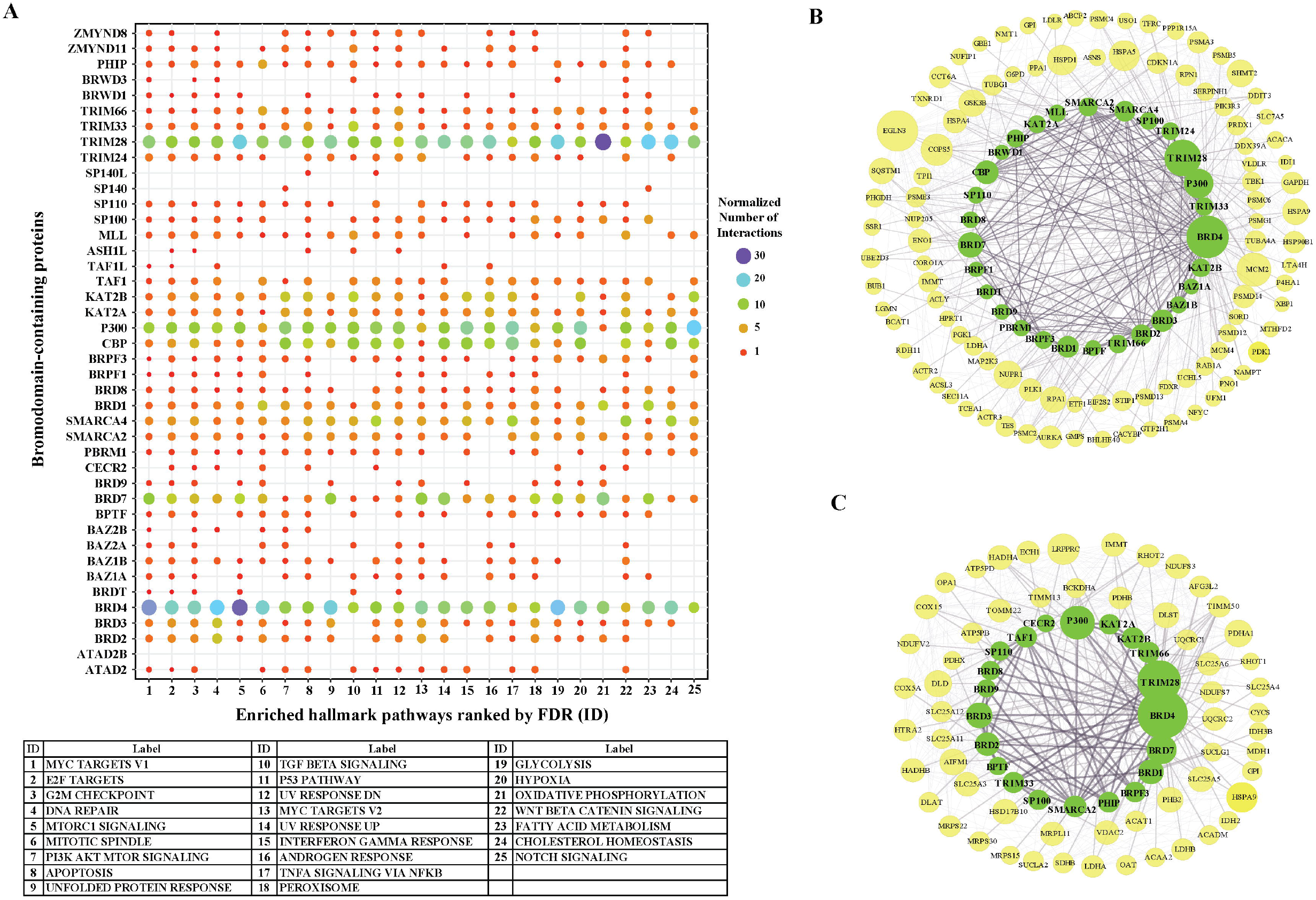

TRIM28 interactions showed the highest enrichment with the Hall-mark oxidative phosphorylation pathway. TRIM28 was previously shown to form a cancer-specific ubi-quitinase together with MAGE-A3/6 proteins (Pineda & Potts, 2015) for proteasomal degradation of AMPK, a master regulator of metabolic/energy homeostasis and mitochondrial biogenesis in cancer cells (Chaube & Bhat, 2016). Consistent with this, enrichment analysis shows a relatively higher significance of TRIM28 interactions associated with glycolysis. 21 BRD proteins also interact with the oxidative phosphorylation pathway, including other TRIM proteins (**Figure 6C**). In this sub-network, BRD4 and BRD7 have relatively more interactions with oxidative phosphorylation path-way members. Collectively, this analysis supports the role TRIM28 and other BRD proteins to modulate cell metabolism and homeostasis by interacting with the protein members in oxidative phosphorylation pathway and related signaling pathways.

We used MTGO (Vella et al., 2018) to further investigate the clustering profile of BRD proteins in BRD PPIN based on gene ontology, as well as the topological features. MTGO clustering identified 70 functional modules, with 16 of them containing BRD proteins (**Table 5**). The cluster containing the most BRD proteins are annotated as transcription factor binding, and this cluster includes 18 BRD proteins: ATAD2, BAZ1A, BAZ1B, BPTF, BRD2, BRD7, BRD9, BRWD1, CBP, P300, KAT2A, KAT2B, MLL, PBRM1, SMARCA4, TAF1, TRIM24, TRIM33. The second largest cluster contains 4 BRD proteins: BRD3, BRD4, BAZ2B and ZMYND11. BRD4 is previously reported to contribute to the regulation of alternative splicing via co-localizing and interacting with the splicing regulators (Uppal et al., 2019). ZMYND11 is reported to regulate RNA splicing via connecting with histone H3.3K36me3 and then interacting with RNA splicing regulators, including the U5 snRNP components of the spliceosome, such as EFTUD2 (Guo et al., 2014). But the roles of BRD3 and BAZ2B in RNA splicing are unclear. There are BRD proteins not included into any the Gene Ontology terms, so their clustering attributes are largely dependent on the interaction profiles.

**Table 5.**
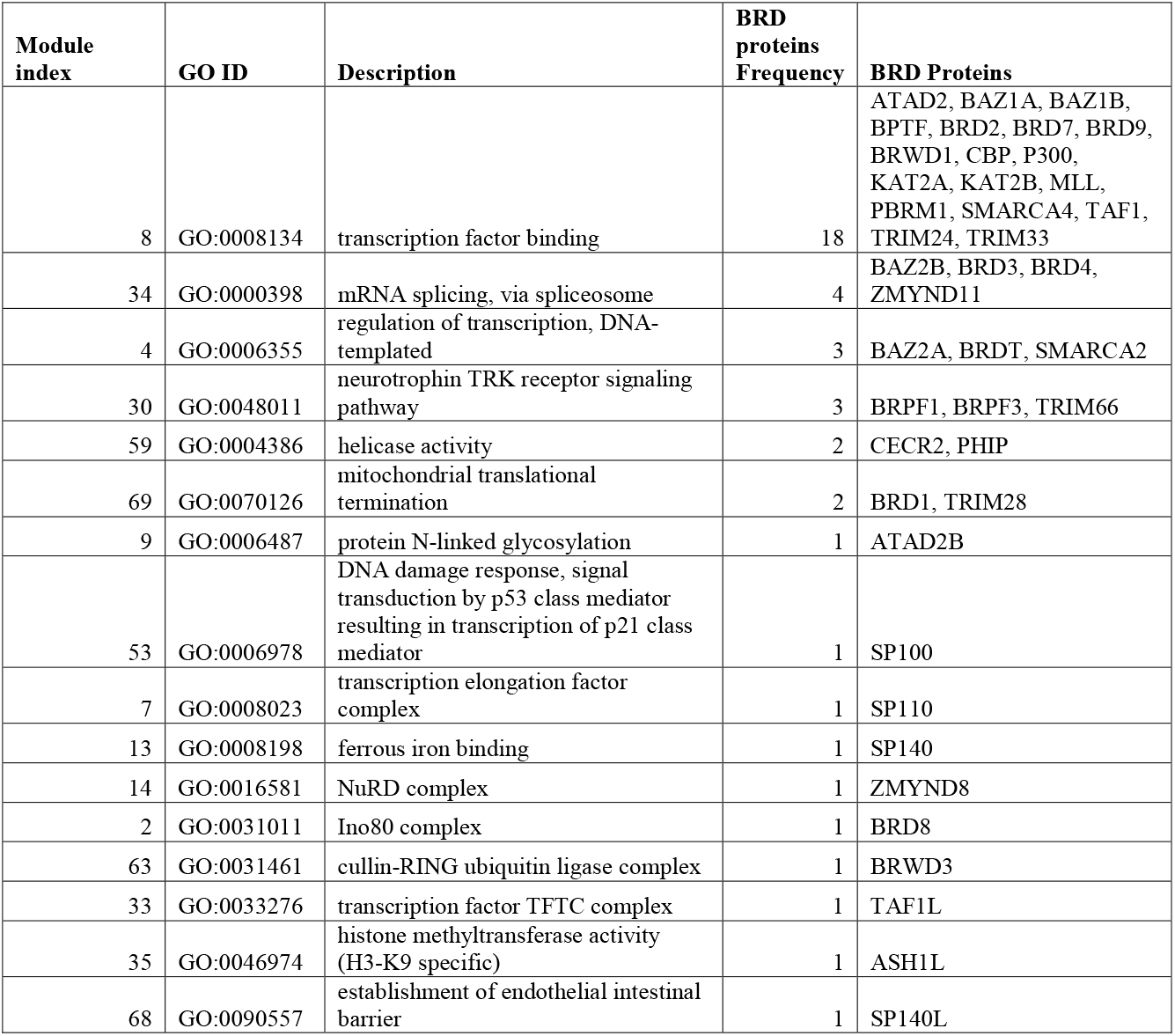
MTGO clustering results for BRD Proteins

Interestingly, some BRD proteins with medium degrees cluster with similar ontological functions, such as BRPF1, BRPF3 and TRIM66 in the neurotrophin TRK receptor signalling pathway, CECR2 and PHIP to RNA helicase activity, and BRD1 and TRIM28 in mitochondrial translational termination via interactions. The clustering results show some potential functions for these poorly studied BRD proteins. Based on MTGO cluster results, ATAD2B is possibly associated with protein N-linked glycosylation. It is unclear if BRD proteins have any role in N-linked glycosylation and the functional role of ATAD2B needs further investigation. Taken together, the interaction network of BRD proteins demonstrates that BRD proteins perform important roles in the context of cellular and biological pathways, and the network topology of BRD proteins provide new insights into their potential functions.

## 4 Discussion

Bromodomain proteins have versatile roles of recognizing acetylated histone and non-histone proteins and forming protein-protein interactions at chromatin to regulate diverse biological processes. Despite a growing number of systematic proteomic studies and a wealth of physical protein interaction data deposited into databases, the role of BRD proteins as mediators of protein complexes in the human PPI network remains unclear. Overall, different BRD proteins have a large number of reported interactions across many studies. Some have a large number of PPIs despite a relatively smaller number of publications. This can be attributed to the use of high-throughput techniques, such as Affinity Purification Mass Spectrometry (AP-MS). For example, TRIM66 interactions were missing in the databases until a single publication in 2018 revealed 200 various interactions for TRIM66 involved in DNA repair response (Kim et al., 2019). Still there is a deficiency of interaction data for a subset BRD proteins, including BRDT, SP140, SP140L and ATAD2B. The cell-specific expression restrictions and/or the relatively low expression levels may contribute to the insufficient studies to these BRD proteins with poorly characterized functions. Thus, several BRD family members remain uncharacterized and systematic analysis of available protein interaction data is needed to predict their functional roles.

In this study, we constructed a network of all BRD proteins and their interactions and applied graph analysis to examine their topological characteristics in the human PPI network. Our systematic analysis provides understanding the global roles of BRD proteins as mediators of protein interactions. Consistent with their role as molecular scaffolds, we found that BRD proteins are hub proteins and form functional protein complexes to shape the human PPI network. Analysis of topological network features support the important organizational roles for BRD proteins in the global protein-protein interaction network. BRD proteins are positioned at the global center, a characteristic that supports their essential and evolutionary conserved functions.

Analysis of the BRD PPI network further highlighted the relationships between BRD proteins. The similarity scores calculated for interactions formed by different BRD proteins provides insight into the shared functions among different BRD proteins, including the less well-characterized BRD proteins. We first determined similarity scores using a distance of 1 and identified four related BRD protein pairs; CBP and P300, SMARCA2 and SMARCA4, KAT2A and KAT2B, BRD2 and BRD3. These BRD proteins all share structural and functional similarities. The cliques formed by CBP, p300 and KAT2B expand to 4-nodes clique with the addition of KAT2A. Since these 4 proteins are all the members of BRD proteins with intrinsic histone acetyltransferase activities, they may work with other non-BRD proteins to form chromatin-modifying complexes, such as the SAGA complex (Soffers & Workman, 2020). CBP and p300 form cliques with other BRD functional group members from other functional clusters, including SMARCA2, SMARCA4 and BRD7. In the 25^th^ maximal clique, CBP interacts with SMARCA2, SMARCA4, TRIM28 and BRD4. CBP/p300 and SMARCA2/4 have been reported to form p300-CBP-p270-SWI/SNF complex (Dallas et al., 1998) to remodel the chromatin structure and thereby regulate gene transcription. In addition, BRD4 interacts with CBP/p300 and SMARCA4 to regulate histone H3 acetylation and chromatin remodeling (Wu, Kamikawa, & Donohoe, 2018). But more study is required to investigate the collective functions of the protein complex predicted by 25^th^ complex.

We modified the similarity score to find similar BRD pairs with distances of 2 and 3, and found that majority of BRD proteins are inter-connected. Several pairs of BRD proteins have common interactions, indicating functional similarity between different BRD protein members. As expected, hub BRD proteins in the global human PPI network exhibit overall higher similarity scores with BRD proteins, partly due to the extensive studies available. BRD proteins are also more likely to have similar interactome profiles with the other BRD proteins falling in the same functional groups. Interestingly, some proteins belong to different functional groups also have higher similarities (greater than 0.5). This indicates BRD proteins from these similar functional groups are related to each other and potentially form protein complexes via common interactors to perform similar biological functions. Examples include the BRD7-CBP-SWI-SNF complex consisting of BRD7, SMARCA2, SMARCA4 and CBP and/or the ALL-1 super complex formed by MLL, TAF1 and SMARCA2 (Giurgiu et al., 2019).

We used enrichment analysis with Hallmark and Gene Ontology pathways to examine the functional roles of BRD protein interactors. Among the top enriched pathways, a large set of BRD proteins (32/42) and their interactions are associated with DNA damage repair responses. The DNA damage repair response (DDR) is carried out by a network of factors that sense DNA damage and signal the recruitment of chromatin remodeling and DNA repair machinery to sites of DNA damage. The BRD proteins are integral to DNA repair responses and participate recognition of acetylation signals, recruiting DDR and transcriptional factors, regulating transcription and remodeling chromatin activities, and triggering DSB repair (reviewed in (Chiu, Gong, & Miller, 2017)). The mammalian SWI/SNF (mSWI/SNF) complexes are ATP dependent chromatin remodeling complexes that contain a bromodomain module and regulates the accessibility of genomic elements for DNA damage repair (Hargreaves & Crabtree, 2011). SMARCA2 (also known as BRM for brahma homologue), SMARCA4 (BRG1, for Brahma-related gene-1), BRD7 and PBRM1 (BAF180) (Kadoch et al., 2013; Kumar, Li, Muller, & Knapp, 2016) are members of these chromatin remodeling complexes. Interestingly, these and 28 other BRD proteins also participate in DDR via interacting with this pathway members (**Figure 5B**). The DNA-repair associated BRD proteins belong to 8 functional groups, with only AAA-ATPase BRD proteins (ATAD2 and ATAD2B) are not encompassed. However, Kim, et al. have suggested ATAD2B is involved in homologous recombination by performing DSB repair assay after knocking down ATAD2B by siRNA (Kim et al., 2019).

Functional characteristics of proteins can be predicted via PPI clusters that share similar interactions, the potential functional roles of poorly studied BRD proteins can be predicted. We therefore performed functional clustering analysis with gene ontology and identified potential functions for poorly characterized BRD proteins. Speckled proteins tend to be expressed in blood cells and have been related to immune cell functions (reviewed in (Fraschilla & Jeffrey, 2020)). As expected, SP100, SP140 and SP140L BRD proteins have been assigned to immune-related clusters in MTGO results (**Table 5**). Similarly, BRD3, BRD4, BAZ2B and ZMYND11 formed a functional cluster of BRD proteins with potential roles in RNA splicing and thus predicted potential functions for less studied BRD proteins. Whether they have functional roles underlying these biological process needs to be further studied.

Advances in systems biology, including an ever-expanding catalog of protein-protein interactions and the development of modern methods for topological and functional prediction have significantly enhanced our ability to study the structure and function of biological networks. In this work, we constructed a PPIN to provide a global view of the protein interactome in humans for the study of the family of BRD proteins. BRD proteins have emerged as central factors in diverse biological processes, yet many BRD proteins remain poorly characterized. Identifying the relationships between BRD protein interactions and functional modules in gene interaction networks is a critical step towards understanding their biological roles. Distinctive hallmark pathways and GO terms were identified in our BRD protein sub-network, and this functional annotation offers new insight for investigation of BRD protein function for both well-studied and unclassified BRD proteins.

Prospective analysis will be useful to exploit the topological and functional modules to define disease modules. A particularly interesting goal is to integrate PPI modules with co-expression networks in specific physiological/pathological contexts. In this way, the comparison of BRD protein functional and topological sets can be compared between disease versus healthy networks to uncover network rewiring events to characterize the detailed events particular disease and to help pinpoint biologically and therapeutically relevant proteins.

## 5 Conflict of Interest

The authors declare no conflicts of interest.

## 6 Author Contributions

Conceptualization: CG, SF; Writing-original draft preparation and literature search: CG, SF; Writing-review and editing: CG, SF, KCG; Supervision: KCG, SF; All authors have read and agreed to the published version of this manuscript.

## 7 Funding

This study was supported by the National Institutes of Health, National Institute of General Medical Sciences and National Cancer Institute award numbers R01GM129338 and P01CA240685, respectively, to KCG and SF.

## 8 Data Availability Statement

R scripts for all analysis are available on GitHub (github.com/frietzelab/BRD_PPIN).

## Notes

### Competing Interest Statement

The authors have declared no competing interest.

