## Supplementary Figures for "Functional networks of the human bromodomain-containing proteins": Supp1_BioGRID.pdf

### Supplementary Figure 1

**A**

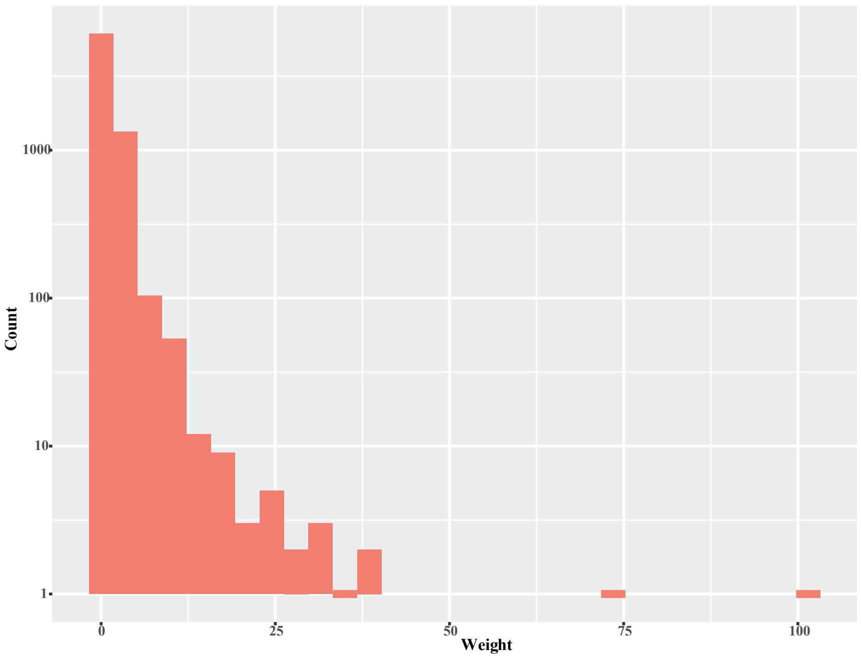


**B**

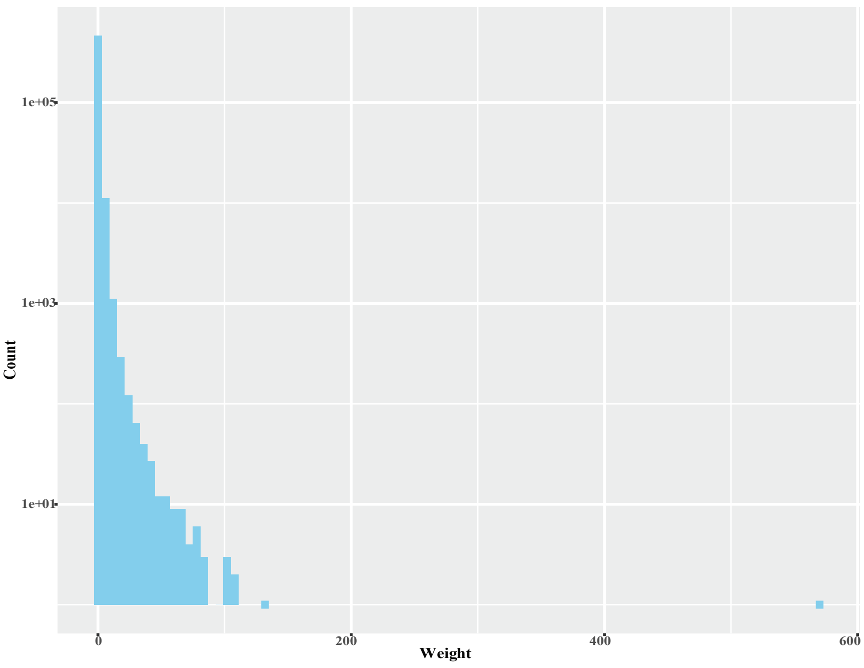


**The distribution of publication/technique number(weight) of interactions in BioGRID.** This figure shows the interaction information for the interactions derived from BioGRID database. X-axis is the number of different techniques and/or publications for interactions; there is case in which different techniques used to confirm the same interaction in the single publication, so the count is the number of techniques. Otherwise, the count refers to the number of publications. Y-axis shows the count of different interactions that have specific numbers of publications/techniques.

A) Publication weights of protein-proteins interactions derived from BioGRID in the global PPIN.

B) Publication weights of protein-proteins interactions from BioGRID in BRD PPIN.
