## Supplementary Figures for "Functional networks of the human bromodomain-containing proteins": Supp2_HIPPIE.pdf

### Supplementary Figure 2

**A**

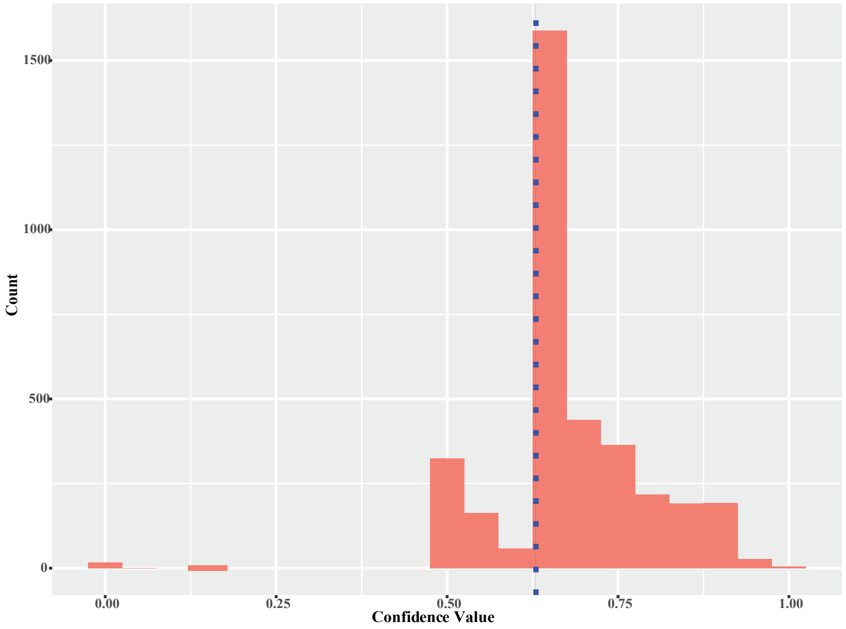


**B**

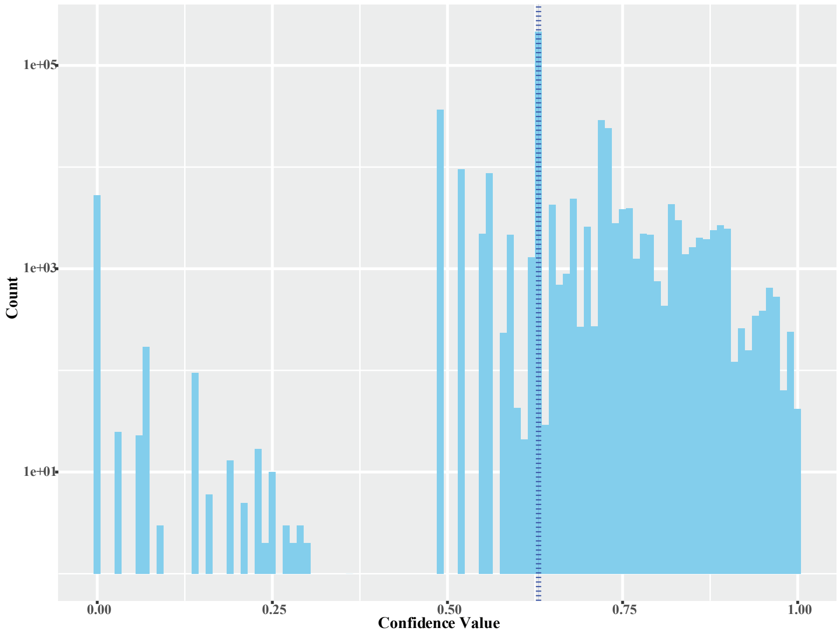


**The distribution of confidence valuse of interactions in HIPPIE.** HIPPIE database got its own algorithm to compute the confidence value of each interaction based on the publications and techniques used, etc. The blue dotted lines refer to the medium confidence value 0.63 suggested by HIPPIE.

A) Confidence values of protein-proteins interactions derived from HIPPIE in the global PPIN.

B) Confidence values of protein-proteins interactions from HIPPIE in BRD PPIN.
