## Supplementary Figures for "Functional networks of the human bromodomain-containing proteins": Supp3_DataSource.pdf

### Supplementary Figure 3

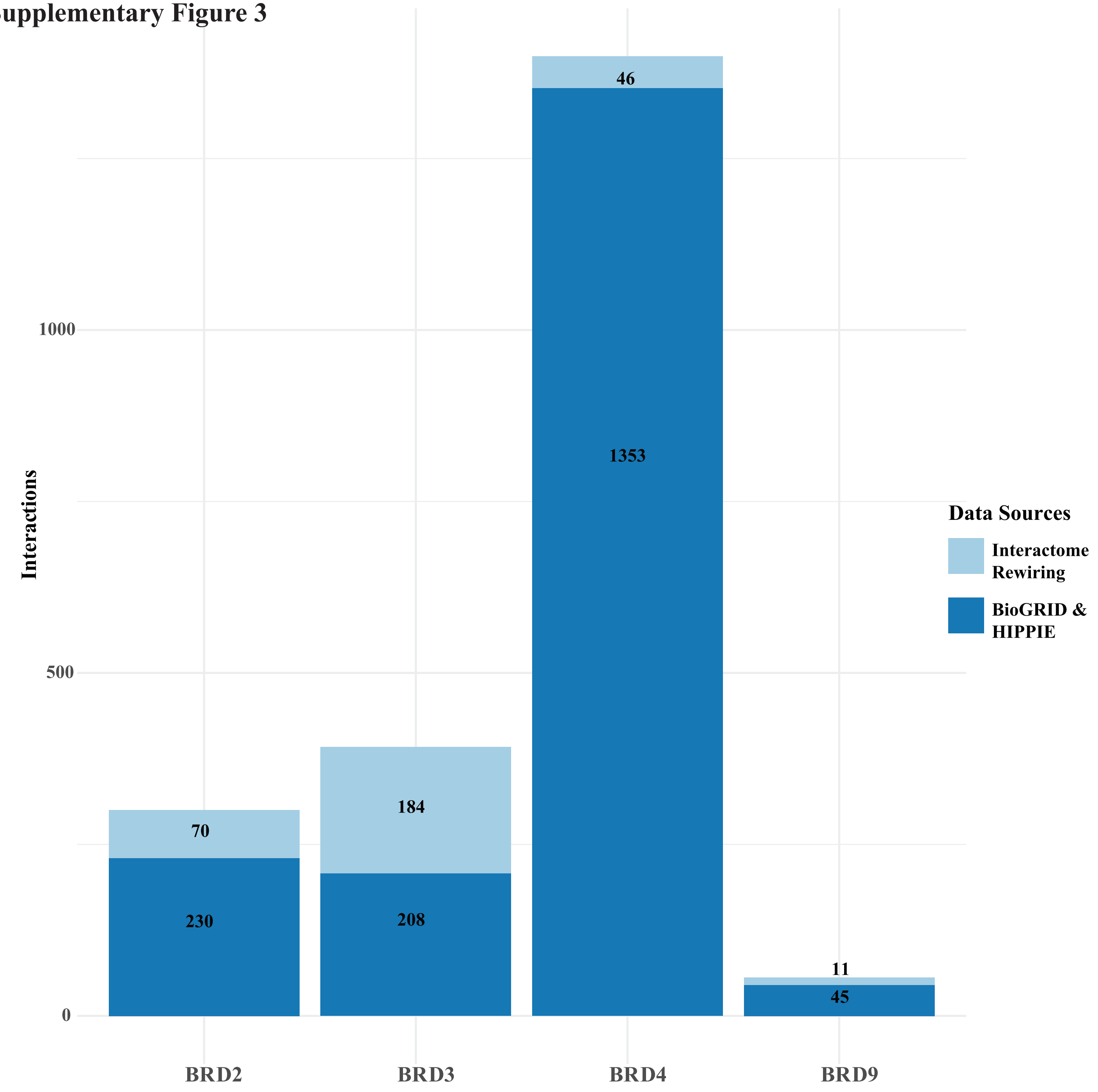


**Interaction summary of BRD2,3,4 and BRD9 derived from public databases and the recent publication.**
X-axis: 4 different BRD proteins; Y-axis: number of interactions from different sources.
