## Supplementary Figures for "Functional networks of the human bromodomain-containing proteins": Supp4_Degree~Pub.pdf

**Supplementary Figure 4**

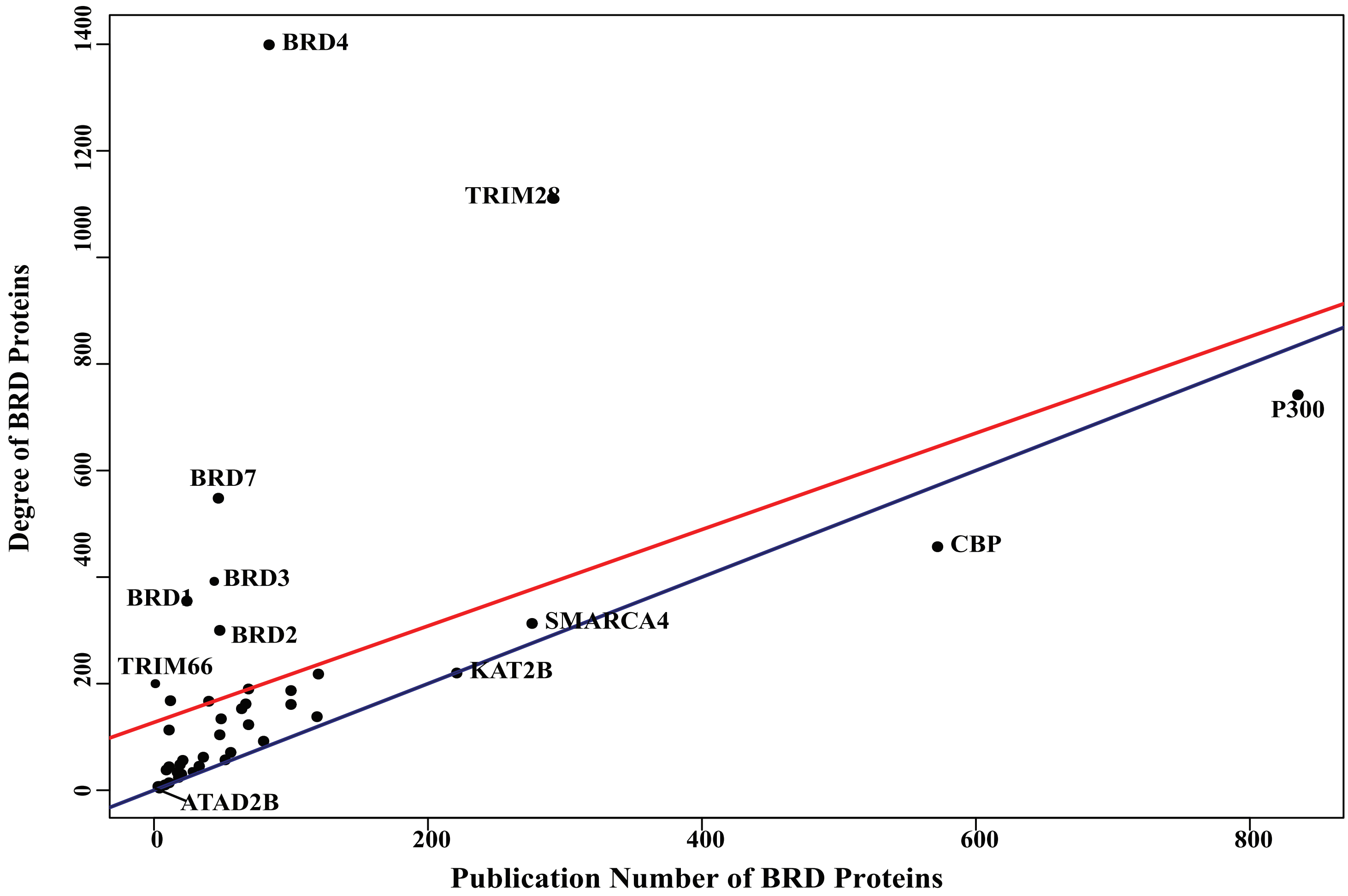


**Relationship between degrees and publications for Bromodomain-containing proteins.**
Relationship between degrees (y-axis) and publication numbers (x-axis) for BRD proteins, the red line is the linear regression line between degree and publication number and the blue line represents a situation where the degree equals the number of publication (a slope of 1). The top 10 hub BRD proteins are labeled, as well as BRD proteins TRIM66 and ATAD2B.
